# Axenisation of oleaginous microalgal cultures *via* anoxic photosensitisation

**DOI:** 10.1101/2024.11.07.622432

**Authors:** A. Iyer, M. Monissen, Q. Ma, M. Osborne, E. Schaedig, O. Modin, R. Halim

**Affiliations:** Conway Institute of Biomolecular and Biomedical Research, UCD, Dublin 4, Ireland; University College Dublin, School of Biosystems and Food Engineering, Dublin 4, Ireland; RWTH Aachen University, Templergraben 55, 52056 Aachen, Germany; Biosciences Center, National Renewable Energy Laboratory (NREL), Golden, Colorado, USA; Architecture and Civil Engineering, Chalmers University of Technology, Gothenburg, Sweden

**Keywords:** Axeny, Rose Bengal, Anoxy, phototrophs, microalgae

## Abstract

Growing interest in sustainable biofuel research has necessitated high quality axenic oleaginous microalgal strains. Unfortunately, most strains available in culture banks contain commensal microbes such as bacteria and the default decontamination method involves antibiotic treatment which has begun to exacerbate the emergence of antibiotic resistance. To overcome this problem, anoxic photosensitisation was investigated as an alternate approach.

Four oleaginous microalgal species (*Tetradesmus obliquus, Desmodesmus armatus, Chlorella vulgaris* and *Nannochloropsis limnetica*) were incubated in varying concentrations of Rose Bengal (0 µM, 1 µM, 3 µM and 9 µM) either in normal (oxic) or anoxic conditions, for 72 h under light (8.85 ± 0.40 W m−2) in a specially designed heterotrophic growth complex (HGC) medium, followed by 72 h in standard Bold’s Basal Medium (BBM). Commonly used antibiotics-based protocol was used as the control method. Post treatment, cell numbers and percentage populations were counted with Flow cytometry, and viability was tested using standard plating methods using BBM and LB. Additionally, the contaminating microbes in the cultures were profiled using 16s rRNA sequencing.

Anoxic conditions were able to significantly decrease bacterial content, albeit with an equally detrimental effect on the microalgal population. Although the responses differed between the microalgae, anoxic incubation along with Rose Bengal at 3 µM was able to completely decontaminate *N. limnetica* and *C. vulgaris*, while *D. armatus* and *T. obliquus* could be decontaminated with an additional streak-plating step. None of the cultures could be decontaminated using antibiotics treatment owing to the presence of gram-negative, multi-drug resistant bacteria such as *S. maltophilia, M. foliorum* and *S. chilensis*.

These results suggest that decontamination of xenic microalgal cultures was largely due to anoxy, that was synergistically enhanced by Rose Bengal at a concentration of ≥3 µM.

**Highlights:**

1. Rose Bengal at 3 µM in anoxic conditions can help achieve axenic microalgal cultures.
2. Standard antibiotics were unable to decontaminate any of the cultures owing to the presence of multi-drug resistant bacteria.
3. Standard antibiotics treatment was deleterious to *D. armatus* cultures.
4. Streak plating may be required for *D. armatus* and *T. obliquus* after anoxic Rose Bengal treatment to obtain individual colonies for further inoculation.

## 1 Introduction

With growing impetus to achieve sustainable climate targets through the generation of sustainable fuels and carbon capture (Goal 7 of U. N. Environment (2017)), microalgal research has not only seen a significant boost in research interest, but more resources are now being channelled towards studying their molecular mechanisms. This has raised the demand on high quality oleaginous strains; many of which are sensitive to contaminating microbes in the culture (Zhu et al., 2020). Many microalgae of interest; though easily available in culture banks, are xenic and contain bacteria or other microbial contaminants that interfere with further upstream research work. Consequently, axenisation / decontamination procedures remains an active topic of research. The most commonly employed first line of treatment to tackle bacterial loads is with antibiotics (Molina et al., 2019) that consequently, has resulted in resistance development owing to indiscriminate use (Peekate & Abu, 2015). While newer methods have been developed with a higher focus on physical separation through microfluidics (Godino et al., 2015; Ota et al., 2019; Syed et al., 2018), and advances in flow cytometry allows for more precise sorting of cells in larger volumes of culture (Scholz, 2014), many of these methods become unreliable when working with bacterial-microalgal associations that are physically attached. Bacteria can lodge on the microalgal surface and become very difficult to remove. Using vigorous physical methods or harsh antibiotic use can become deleterious to the microalgae itself. Moreover, while many protocols call for a combination of chemical and physical methods to attain axenic cultures though such procedures quickly become protracted, expensive, and can significantly hamper the survivability of the microalgal culture.

The work described in this publication attempts to investigate the combined physico-chemical treatment on contaminated microalgal cultures through anoxy and photosensitisation to achieve axenic cultures. Photosensitisation through the use Rose Bengal appears as a lucrative method of decontamination against bacterial and viral particles (Cossu et al., 2016); though the concentrations noted in the publication were in the mM range. Bacteria generally do not posses the necessary pathways to mitigate the production of free oxygen radicals that are generated when Rose Bengal is exposed to light in an aqueous medium (Lambert et al., 1990). Moreover, being a xanthene derivative, Rose Bengal posses anti-bacterial properties even in the absence of light and has been used in formulation with chloramphenicol to isolate fungi from mixed cultures (Ottow, 1972). Likewise, removal of oxygen from cultures has been observed to greatly aid decontamination procedures (Ma et al., 2017). While this is generally achieved through flushing the growing microalgal culture with CO_2_ or N_2_., the pH requires regular adjustment.

The experimental setup described herein sought was to leverage the ability of microalgae to not only survive under anoxic conditions through the production of its own oxygen via photosynthesis, but also utilise its photosynthetic apparatus to expunge excess oxygen radicals generated from the Rose Bengal dye and selectively culture them free of other contaminating organisms.

## 2 Materials and methods

Water used in the experiments was purified via a Milli-Q^®^ purification system. The experiment was performed in biological triplicate. Sterile techniques were performed in a Laminar flow hood (Hearus, Ireland) with all equipment doused in 70 % (v/v) denatured alcohol, and exposed to UV light for at least 10 min prior to commencement of experiments. Anaerobic / anoxic conditions were maintained in an anaerobic chamber (Whitney Biosciences).

Dye Stock Solution (DSS, 10 mM, 10 mL) was prepared using Rose Bengal Dye (101.7 mg, Thermo Fisher Scientific, India) and sterilised by filtering through a sterile 0.22 µm filter (Millex GV, PVDF membrane).

### 2.1 Media

Incubation of microalgae in Rose Bengal was carried out in a heterotrophic growth complex (HGC, 200 mL, pH 7.2): Tryptone (10 g L^−1^; GBiosciences; St. Louis; USA), Yeast extract (5 g L^−1^; UltraPure; Thermo Scientific, USA), Coral Pro Salt (1 g L^−1^; Red Sea, Israel), and pyruvic acid (20 µg L^−1^; Merck, Germany) to supplement growth even during the continuous dark phases. For anoxic conditions, 100 mL medium was autoclaved and then placed in the anaerobic chamber and allowed to equilibrate under constant stirring for five days. For aerobic incubation conditions, 100 mL of the medium was autoclaved and used without any further modifications.

Mother cultures and post-treatment culturing was carried out in standard 3N+BBM (Culture Collection of Algae and Protozoa, 2019). Similar to HGC, anaerobic 3N+BBM (100 mL) was prepared by autoclaving the medium and placing it in the anaerobic chamber for five days and allowing equilibration for five days under constant stirring.

### 2.2 Microalgal species

Xenic mother cultures (900 mL) of *Desmodesmus armatus, Tetradesmus obliquus, Chlorella vulgaris*, and *Nannochloropsis limnetica* cultures were maintained in ambient (21 °C to 25 °C) temperature conditions under constant shaking (80 rpm) with a 14 h light cycle at 8.85 ± 0.40 W m^−2^ brightness (Barrina T5 LED Grow Lights, Full spectrum, USA).

### 2.3 Anoxic treatment conditions

Under aseptic conditions in a laminar hood, the xenic microalgal mother culture (50 mL) was drawn in sterile centrifuge tubes (Falcon^®^, BD), centrifuged at 3000 × *g* for 10 min at 15 °C. The supernatant was aseptically discarded and the cells were brought into the anaerobic chamber and allowed to equilibrate for 5 min to facilitate air exchange in the tube’s head space. Anaerobic HGC medium (12 mL) was then added to the tube and the cells were gently resuspended through repeated aspiration.

The microalgal cells suspended in anaerobic HGC (1 mL) was added to the 4 mL HGC medium with experimental Rose Bengal concentrations and the culture tubes were screwed shut, and sealed by wrapping Parafilm^®^. The tubes were then taken out of the anaerobic chamber and incubated on a roller-mixer for 72 h under 14 h light cycles with ambient temperature conditions (21 °C to 25 °C).

The samples were then centrifuged at 3000 × *g* for 10 min at 15 °C and the tubes were taken into the anaerobic chamber. There, the supernatant was discarded and the cells were gently resuspended in anaerobic 3N+BBM (5 mL each) through repeated aspiration. The tubes were sealed once again, taken out of the anaerobic chamber, and incubated on a roller-mixer for 48 h under 14 h light cycles with ambient temperature conditions (21 °C to 25 °C).

### 2.4 Oxic treatment conditions

Oxic treatment of the microalgae was identical with the anoxic treatment noted above albeit without the use of the anaerobic chamber. All sterile transfers and inoculations were carried out under the laminar hood.

### 2.5 Antibiotic treatment conditions

Antibiotic treatment was carried out as a comparator / control condition. The method used was described previously by Han et al. (2016), with a few modifications. Briefly, a mixture of Ampicillin, Gentamycin sulfate, Kanamycin, Neomycin and Streptomycin (600 mg L^−1^ each) in 3N+BBM was prepared and sterile filtered using 0.22 µm filter (Millex GV, PVDF membrane). Microalgal cultures (5 mL) was centrifuged at 3000 × *g* for 10 min at 15 °C and the supernatant was discarded. The microalgae was resuspended in the antibiotics mix (5 mL) and suspended for three days and then transferred to antibiotics-free BBM for five days.

### 2.6 Characterisation of contaminants

The contaminating microbes in the mother culture were profiled using 16s ribosomal RNA sequencing. The forward primer Bac27F (5^′^-AGAGTTTGATCCTGGCTCAG-3^′^) and universal reverse primer: Univ1392R (5^′^-GGTTACCTTGTTACGACTT-3^′^) were used. For PCR, initial denaturation was at 95 °C for 5 min; 35 cycles of denaturation (30 s at 94 °C), annealing (30 s at 55 °C), and extension (2 min at 72 °C); and a final extension at 72 °C for 7 min (Jiang et al., 2006). The PCR-amplified products were purified using a subjected to Qiagen clean-up kit QIAquick PCT Purification (CAT:28106) and sequenced at Eurofins using the Eurofins Genomics Mix2Seq platform with Bac27F and Univ1392R primers. BLAST (Altschul, 1997) was used to compare the sequences to the 16S ribosomal RNA sequences database in NCBI GenBank database (Sayers et al., 2022).

### 2.7 Tests for axeny and viability

Axeny was deduced by considering the results from three tests:

#### 2.7.1 LB plating

Sterile petri dishes with 10 mL of Lysogeny Broth Agar (Lennox L Agar, Invitrogen^(tm)^, USA) were prepared. The dishes were divided into three sections and the culture sample (10 µL) was spread in each section. The plates were incubated at 30 °C for five days to check for any growth. If growth was detected, the colony was picked and observed under light microscopy.

#### 2.7.2 BBM plating

BBM plates with agar (2 % w/v) were prepared and 10 µL of sample was spread on the plate and incubated in ambient conditions for five days to check for any growth. If growth was detected, the colony was picked and observed under light microscopy.

#### 2.7.3 Flow cytometry

##### Samples

Samples (1 mL) from the mother culture and post treatment culture growth were analysed using a CytoFLEX LX flow cytometer (Beckman Coulter, USA) to compare difference in bacterial contamination. From each 1 mL sample, two 250 µL aliquots were drawn. One aliquot was for staining and the other was the unstained control.

##### Cell fixing

Each aliquot (250 µL) was added to paraformaldehyde (4 % v/v, 250 µL) and mixed for 10 s in a vortex mixer, and allowed to rest 10 min at 4 °C. The sample was centrifuged at 3000 × *g* for 5 min at 4 °C and part of the supernatant (450 µL) was discarded and replaced with 100 µL of paraformaldehyde and mixed again for 5 s using a vortex mixer.

##### Cell staining

To each fixed aliquot (250 µL), SYBR Green I (28 µL, 1:1000 dilution in dimethyl sulphoxide) was added and vortexed for 10 s and allowed to rest in the dark for 15 min. SYBR Green I was obtained from Invitrogen (Thermo Fisher Scientific, USA).

##### Measurement

Autofluorescence was observed using Cytoflex LX and under FSC-A vs R660-APC for chlorophyll and non-chlorophyll pigments. Likewise, FSC-A vs B525-FITC-A was observed for fluorescence from SYBR Green that stained the bacterial DNA, that was distinguished from microalgal cells by size and autofluorescence.

### 2.8 Final axenic culture

Final purification step was performed using 50 µL samples under the 3 µM anoxic Rose Bengal concentration on BBM agar using the standard four quadrant streak-plating method (Sanders, 2012) and incubated for 120 h. No antibiotics were included in the medium. Single colonies from the final quadrant were picked and cultured in 3N+BBM under the conditions mentioned in subsection 2.2. After 36 h of culturing, samples (1 mL) was tested for 16s rRNA amplification and streaked on LB plates to check for bacterial growth. Microscope images of the final culture was taken and compared to the mother culture.

### 2.9 Statistical analyses

All statistical analyses were preformed using R (4.4.0) (R Core Team, 2024) with RStudio as the frontend (RStudio 2024.04.1+748). Data wrangling and analyses were carried out using the tidyverse set of packages (Wickham et al., 2019). Flowcytometry data was collected using the proprietary CytExpert software (Version 2.6, Beckman Coulter, USA) bundled along with the CytoFLEX LX (Beckman Coulter, USA), but subsequent analyses was performed using FCS Express software (Version 6.06, Dotmatics, USA).

## 3 Results

### 3.1 Quantitative assessments of axeny and viability

The percent proportions of microalgae and bacterial cells in the culture were quantified using flow cytometry, and the values are provided in Table 1.

**Table 1:** Quantitative scoring for presence of microalgae and other microbes in culture using light microscopy.

| RB | Oxy | *N. limnetica* culture |  | *D. armatus* culture |  | *C. vulgaris* culture |  | *T. obliquus* culture |  |
| --- | --- | --- | --- | --- | --- | --- | --- | --- | --- |
|  |  | Microalgae | Others | Microalgae | Others | Microalgae | Others | Microalgae | Others |
| 9 | Oxic | 76.9±4.05 | 29.82±1.33 | 3.40±0.54 | 88.06±1.39 | 25.95±0.55 | 5.49±0.20 | 15.75±0.44 | 75.33±1.54 |
|  | Anox | 2.24±0.55 | 0.16±0.01 | 0.05±0.04 | 0.17±0.00 | 1.64±0.88 | 0.37±0.06 | 5.78±0.99 | 0.65±0.20 |
| 3 | Oxic | 52.74±0.63 | 45.00±0.44 | 5.47±0.95 | 89.78±0.40 | 20.26±0.02 | 3.51±0.00 | 23.74±0.06 | 85.42±2.56 |
|  | Anox | 4.09±0.88 | 0.40±0.47 | 0.12±0.00 | 0.01±0.00 | 5.49±0.06 | 0.57±0.89 | 17.71±0.72 | 0.34±0.08 |
| 1 | Oxic | 59.9±1.30 | 38.66±0.77 | 4.30±0.46 | 90.91±4.12 | 18.56±5.09 | 22.60±0.17 | 24.63±1.11 | 84.72±1.60 |
|  | Anox | 5.1±0.13 | 0.69±0.16 | 0.06±0.00 | 0.33±0.03 | 7.18±1.00 | 0.83±0.22 | 12.58±3.65 | 1.88±0.99 |
| 0 | Oxic | 42.81±1.41 | 43.81±18.3 | 6.39±4.19 | 90.27±5.09 | 12.95±6.03 | 55.55±3.61 | 14.76±5.93 | 79.66±0.38 |
|  | Anox | 7.66±0.18 | 0.66±0.03 | 0.04±0.00 | 0.23±0.2 | 5.87±0.05 | 1.07±0.12 | 4.33±0.09 | 0.67±0.20 |
| Antibiotics |  | 12.5±1.55 | 10.10±3.21 | 8.22±0.0 | 82.34±1.95 | 25.15±6.00 | 5.09±1.06 | 6.98±1.09 | 51.99±2.31 |

Values expressed in % total counts.
Oxy = Oxygen conditions of “oxic” (normal oxygen) or “anox” (low oxygen).
RB = Rose Bengal (μM).
Others = contaminating microbes in the culture.

In all cases, cell percentage fractions of microalgae and bacteria were lower in anoxic conditions compared the ambient oxic conditions.

#### 3.1.1 Analyses of *N. limnetica* culture

##### Oxic conditions

A one-way ANOVA across the experimental conditions for the microalgae *N. limnetica* in oxic conditions, showed significant difference across the groups (F_(4,10)_=374, p=7.7 × 10^−11^), and a pairwise analysis using Tukey HSD found significant difference in the microalgal population values across all the experimental conditions. For the contaminants, there was significant difference across the groups as well (F_(4,10)_=8.91, p=2.48 × 10^−3^) although a closer inspection using Tukey HSD found that only Antibiotics v/s RB 0 (p=0.004), Antibiotics v/s RB 1 (p=0.012), and Antibiotics v/s RB 3 (p=0.003) showed significant difference in population numbers. This suggests that under oxic conditions, the microalgal population undergo change when exposed to varying Rose Bengal concentrations, while the bacterial concentrations only appear to be statistically different between the control antibiotics treatment and the Rose Bengal concentrations, but not within the experimental Rose Bengal concentrations.

##### Anoxic conditions

The ANOVA across the experimental anoxic conditions found an overall significant difference in the *N. limnetica* population (F_(4,10)_=67.47, p=3.39 × 10^−7^), and pairwise Tukey comparison found similarities between RB 9 v/s RB 3 (p=0.12), RB 3 v/s RB 1 (p=0.60), although RB 9 v/s RB 1 was statistically different at 95 % confidence (p=0.013). When considering changes in the bacterial populations, ANOVA was significant overall (F_(4,10)_=26.39, p=2.7 × 10^−5^), and Tukey HSD found similar trends to the oxic conditions where where no significant difference was found between RB 9 v/s RB 3 v/s RB 1 v/s RB 0 (p>0.9), but the significance was largely driven by the control antibiotic condition versus the Rose Bengal conditions (p<0.05).

#### 3.1.2 Analyses of *D. armatus* culture

##### Oxic conditions

A one-way ANOVA across the experimental conditions for *D. armatus* in oxic conditions, showed no significant difference across the groups (F_(4,10)_=2.774, p=0.87). This suggests that the chosen Rose Bengal concentrations do not appear to have an effect on the microalgal populations. In the case of bacterial population, there was some statistical significance noted (F_(4,10)_ = 3.726, p=0.042), but further Tukey HSD only found marginal statistical difference between Antibiotics v/s RB 1 (p=0.045). This suggests that across the treatment conditions under oxic conditions, very little change could be elicited in the D. armatus and its contaminant bacterial population, either through Rose Bengal, or antibiotics.

##### Anoxic conditions

ANOVA analysis of the D. armatus population under anoxic condition found an overall difference (F_(4,10)_=120710, p<2 × 10^−16^) and pairwise Tukey HSD found significant difference between RB 9 v/s RB 3 (p=0.006), RB 3 v/s RB 1 (p=0.016), RB 3 v/s RB 0 (p=0.002), and between Antibiotics and the Rose Bengal conditions (p<0.001). Similarly for the bacterial population, ANOVA found an overall significant difference in the populations (F_(4,10)_=5266, p=2 × 10^−16^), and the Tukey HSD revealed that this was due the differences between the Antibiotics treatment and Rose Bengal concentrations (p<0.001), but not within the experimental Rose Bengal concentrations (p>0.9).

#### 3.1.3 Analyses of *C. vulgaris* culture

##### Oxic conditions

A one-way ANOVA of *C. vulgaris* population across the experimental conditions was found to be significant overall (F_(4,10)_=4.32, p=0.028), and pairwise HSD found significant difference between RB 9 v/s RB 0 (p=0.03) and RB 0 v/s Antibiotics (p=0.043). In the case of contaminating bacteria, ANOVA found significant difference (F_(4,10)_=517.3, p=1.54 × 10^−11^), and Tukey HSD *post hoc* found significant pairwise difference across all conditions except RB 9 v/s RB 3, RB 3 v/s Antibiotics, and RB 0 v/s Antibiotics (p>0.5). This suggests that bacterial content changes significantly, but become comparable with antibiotics use at higher Rose Bengal concentrations.

##### Anoxic conditions

Under anoxic conditions, ANOVA found significant difference across the groups (F_(4,10)_=34.94, p=7.52 × 10^−6^) and a Tukey HSD found that the significance was driven largely by the difference between Antibiotics v/s Rose Bengal conditions (p>0.001). This trend was mirrored for bacterial contaminating cells as well with the overall ANOVA being significant (F_(4,10)_=29.57, p=1.62 × 10^−5^) and Tukey HSD revealing difference between Antibiotics and Rose Bengal conditions (p<0.001). These results suggest that under anoxic conditions, differences between the antibiotics treatment and Rose Bengal treatments are significantly different, but the changes within the experimental Rose Bengal concentrations is minimal.

#### 3.1.4 Analyses of *T. obliquus* culture

##### Oxic conditions

Under oxic conditions, *T. obliquus* showed an overall significant difference in the microalgal populations across the test groups (F_(4,10)_=20.89, p=7.62 × 10^−5^) with Tukey HSD revealing significant differences across all the pairwise comparisons except RB 9 v/s RB 0 (p=0.99), and RB 3 v/s RB 1 (p=0.99). In the case of bacterial contaminants, the trends mirrored the microalgal populations, with ANOVA finding significant overall differences across the groups (F_(4,10)_=166.6, p=4.19 × 10^−9^). All conditions showing significantly different bacterial populations except RB 3 v/s RB 1 (p=0.99), and RB 9 v/s RB 0 (p=0.09). This suggests that under oxic conditions, the effect of Rose Bengal at RB 1 and RB 3 are similar and show the highest microalgal population.

##### Anoxic conditions

A one-way ANOVA of the microalgal populations under anoxic conditions found significant overall difference (F_(4,10)_=29.01, p=1.76 × 10^−5^) with Tukey HSD revealing similar microalgal populations in conditions of RB 9 v/s RB 0 (p=0.853), Antibiotics v/s RB 0 (p=0.417), and Antibiotics v/s RB 9 (p=0.917). For bacterial populations, while the ANOVA found significant overall differences between the conditions (F_(4,10)_=1226, p=2.09 × 10^−13^), the significance was largely driven by the difference between the antibiotics treatment and the Rose Bengal concentrations (p<0.001), but not between the experimental Rose Bengal concentrations (p>0.5). This suggests that at concentrations of 1 µM and 3 µM Rose Bengal, microalgal populations undergo change, but the overall bacterial population numbers remain statistically unchanged unless treated with antibiotics.

Based on these values, anoxic conditions were able to decrease the contaminant numbers, and its effect was somewhat amplified by the Rose Bengal depending on the experimental species. To help visualise the performance of the treatments and decide on a suitable condition, a ratio of contaminant bacteria versus microalgae was calculated and plotted, as shown in Figure 1. Values lower than 1 indicated a lower load of contaminant bacteria relative to the microalgae (i.e. a cleaner culture).

**Figure 1:**
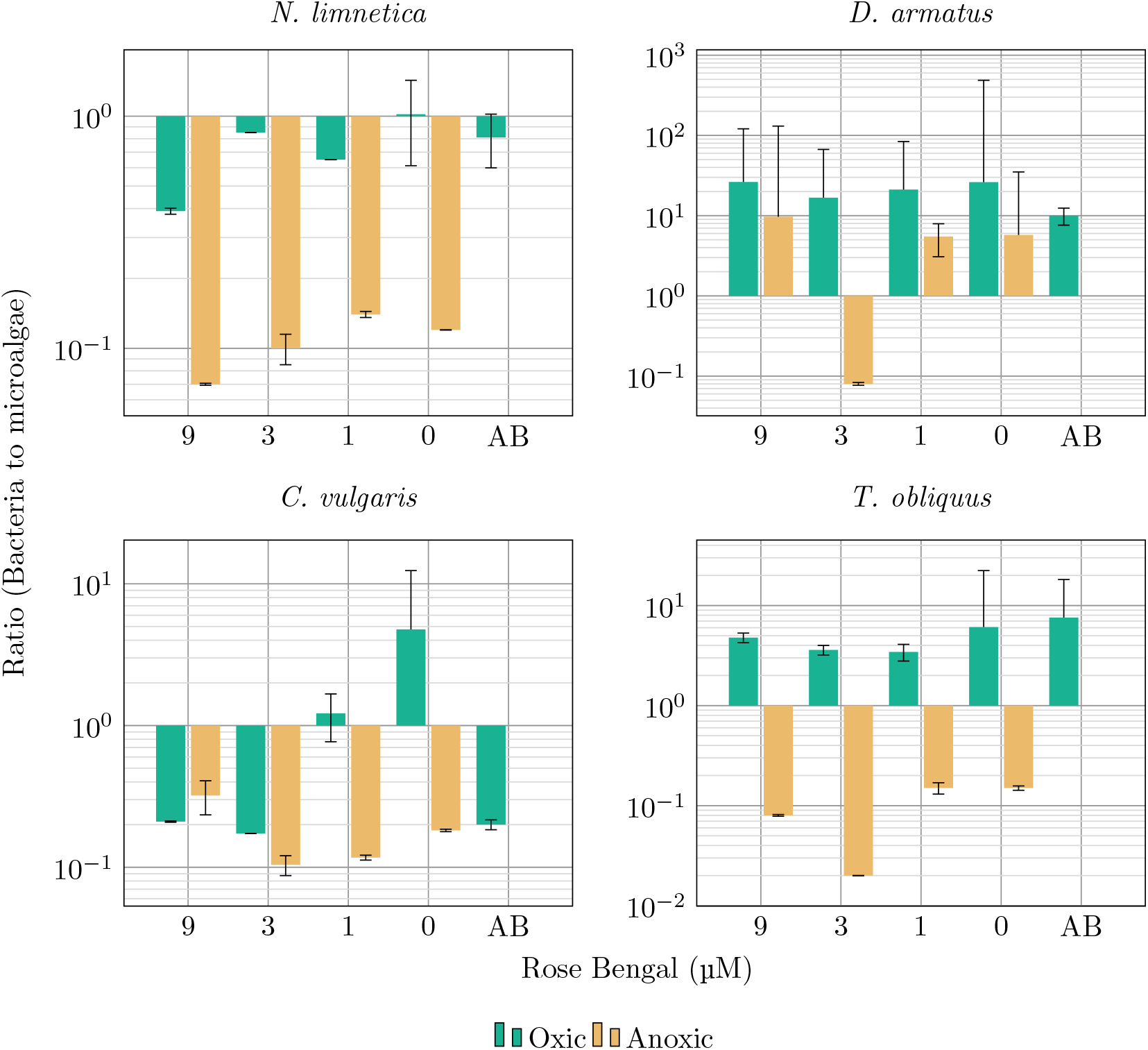
Ratio of contaminant microbes to microalgae in each treatment condition. Lower the value, the higher the number of microalgal cells relative to the contaminant microbes.

Based on Figure 1, favourable bacteria to microalgae ratios were obtained at RB 3 µM for *T. obliquus* and *D. armatus*, while the response ratio was relatively indistinguishable between the Rose Bengal concentrations for *C. vulgaris* and *N. limnetica*. However, when considering the percentage microalgal values from Table 1, anoxic conditions with Rose Bengal concentration of 3 µM appears to be universally effective at maintaining sufficient microalgal population while having a relatively low bacterial content.

### 3.2 Qualitative assessments of axeny and viability

While the cell numbers in Table 1 showed percentage proportions of cell population, it does not attest to their viability, which was checked with BBM and LB agar plating as shown in Table 2 below.

**Table 2:**
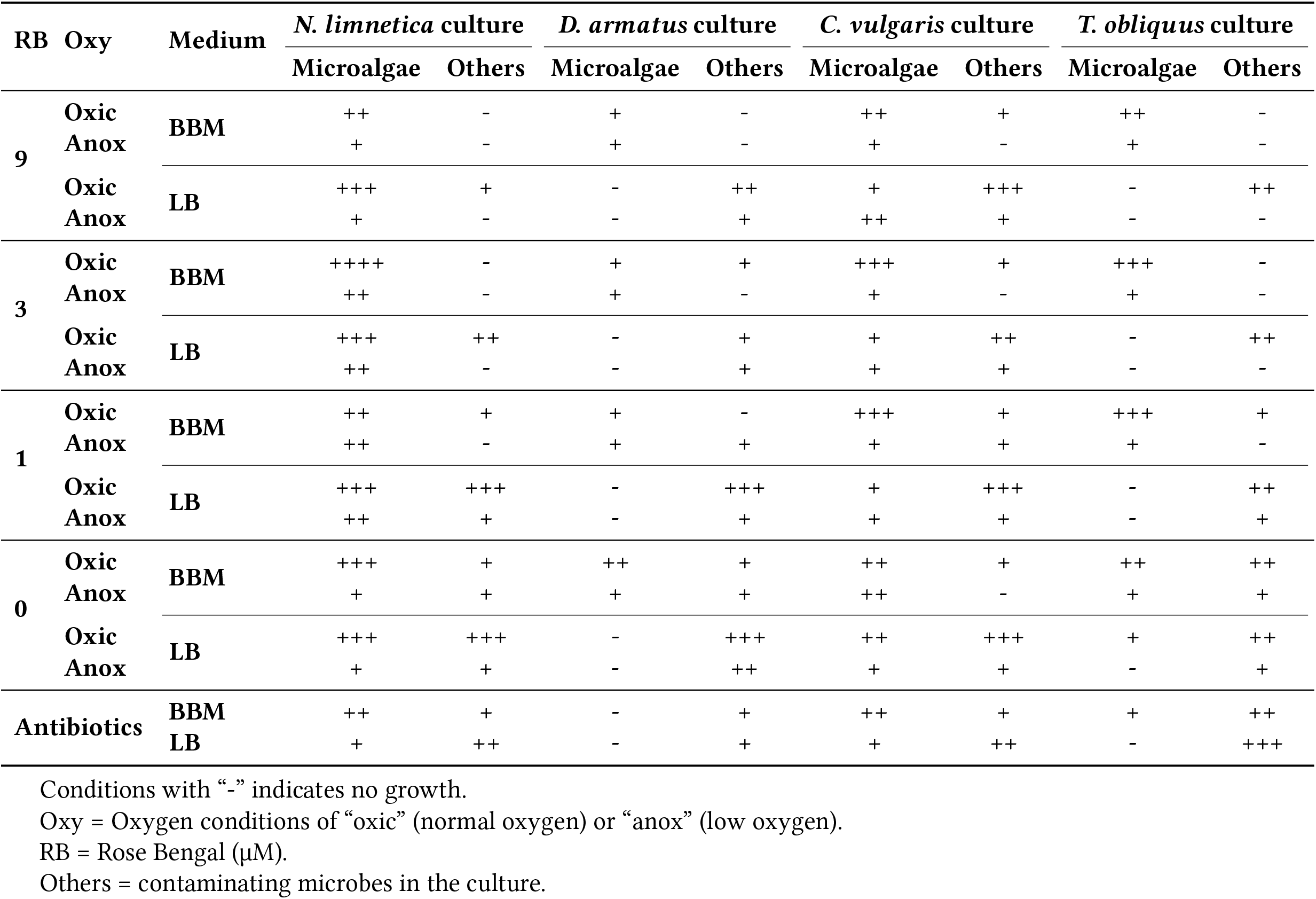
Quantitative scoring for presence of microalgae and other microbes in culture using light microscopy.

The BBM plating was performed to check the growth of the microalgae and the LB agar was used to check bacterial growth. However, it was found that *D. armatus* and *T. obliquus* were incapable of growing on LB agar, while *C. vulgaris* and *N. limnetica* were able to grow in these heterotrophic conditions. Distinction between the colonies however was trivial owing to the exclusive presence of chlorophyll in microalgal cells.

#### 3.2.1 Plating profile of *N. limnetica*

In the BBM plating, it was found that *N. limnetica* growth was the highest at 3 µM Rose Bengal oxic conditions concentration and was the lowest in Antibiotic treatment, 0 µM and 9 µM anoxic Rose Bengal conditions. Bacterial colonies were only observed 1 µM oxic, 0 µM oxic and Antibiotic conditions. This suggests that across all the experimental conditions, viable *N. limnetica* cells were present in the culture.

In the LB plating, bacteria was absent in 3 µM and 9 µM anoxic Rose Bengal concentrations but was detected in all other conditions.

#### 3.2.2 Plating profile for *D. armatus*

In the BBM plating, *D. armatus* was unviable after antibiotic treatment and showed relatively higher growth in 0 µM oxic Rose Bengal treatment. Bacterial growth was observed in 0 µM oxic and anoxic Rose Bengal concentration, 1 µM anoxic Rose Bengal concentration, and 3 µM oxic Rose Bengal concentration. In the LB plating, all conditions showed bacterial colonies although they were the fewest in 3 µM and 9 µM anoxic Rose Bengal conditions.

#### 3.2.3 Plating profile for *C. vulgaris*

In BBM plating, *C. vulgaris* was able to grow well in at Rose Bengal concentrations of 1 µM and 3 µM in oxic conditions. Bacterial colonies were observed in all oxic conditions, but were absent in all anoxic conditions except at 1 µM concentration.

In LB plating, bacteria showed higher growth in oxic conditions compared to anoxic conditions.

#### 3.2.4 Plating profile of *T. obliquus*

In BBM plating, *T. obliquus* showed similar trends to *C. vulgaris* and grew best at Rose Bengal concentrations of 1 µM and 3 µM in oxic conditions.

Similar to *D. armatus, T. obliquus* was unable to grow well in LB conditions and served as a good indicator of exclusive bacterial growth. No bacterial growth was observed in anoxic Rose Bengal concentrations of 3 µM and 9 µM.

### 3.3 Axenic colonies

Based on the results from Tables 1, and 2, streak-plating was performed using cultures treated with anoxic Rose Bengal at 3 µM. Individual colonies were picked to obtain cells that were subsequently inoculated in BBM to obtain axenic cultures. These cultures were then subjected to 16s sequencing using the universal primers, but no amplification was observed. Moreover, subsequent LB plating did not find any bacterial colonies (data not shown). Finally, the cultures were manually inspected under a light microscope. Photographs of the microscope images of the culture prior to, and after treatment is shown in Figure 2 below.

**Figure 2:**
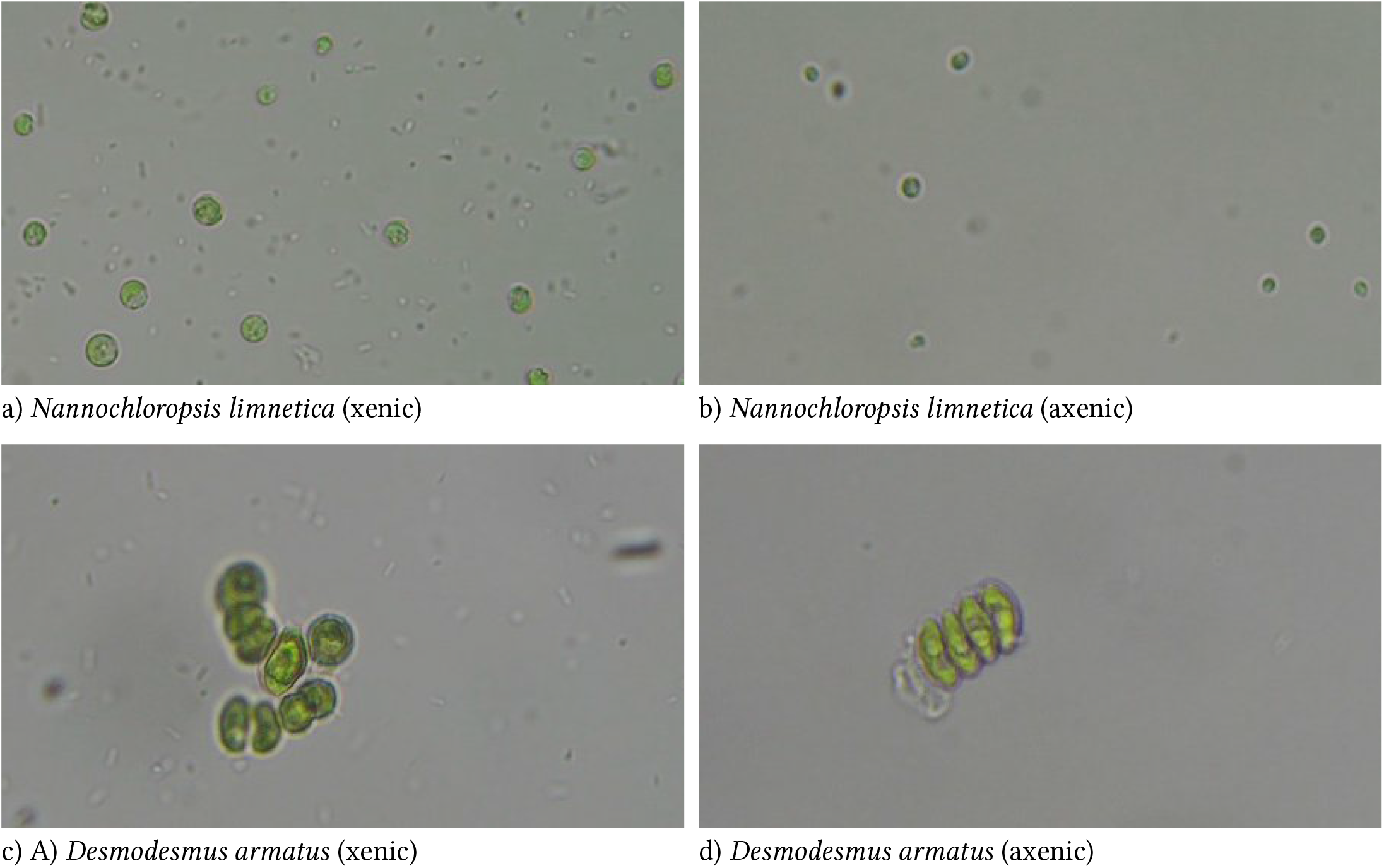
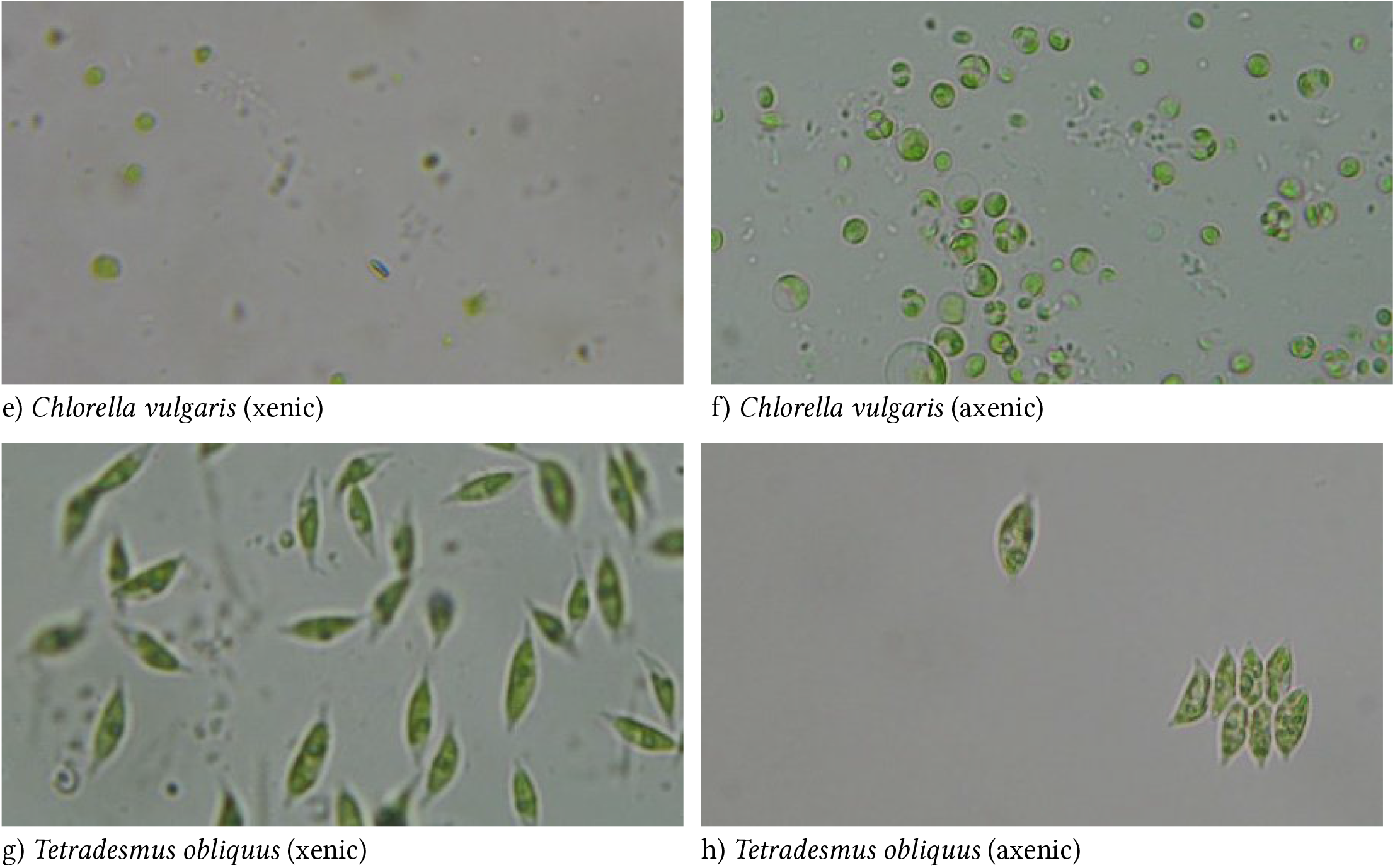
Comparative images of the microalgal cultures before (A, C, E, and G) and after anoxic 3 µM Rose Bengal treatment (B, D, F, and H) of *Nannochloropsis limnetica, Desmodesmus armatus, Chlorella vulgaris*, and *Tetradesmus obliquus* respectively.

In Figure 2 A, C, E, and G, xenic mother cultures of *Nannochloropsis limnetica, Desmodesmus armatus, Chlorella vulgaris*, and *Tetradesmus obliquus* respectively show heavy background of contaminant bacteria while the post treatment samples (B, D, F, and H) show a cleaner profile without any visible bacterial cells.

### 3.4 Mother culture contaminants

The following contaminants were identified in the mother cultures:

A *Stenotrophomonas* sp. contaminant was identified in the *D. armatus, T. obliquus*, and *N. limnetica* cultures, likely *Stenotrophomonas maltophilia*; a multi-drug resistant, gram-negative, opportunistic pathogen (Brooke, 2012). Similarly a member of the *Sphingopyxis* sp; likely, *Sphingopyxis chilensis* was also noted in the *D. armatus* and *T. obliquus* cultures. Members of this genus are capable of degrading chlorophenols and have been proposed for bio-remediation. While they are naturally found along natural water-bodies, they are particularly abundant in soils and stagnant water contaminated with pesticides, fertilisers, and other industrial compounds (Sharma et al., 2021). *S. chilensis* in particular is reported to produce polyhydroxyalkanoate (bio-plastics) (Godoy et al., 2003).

A *Microbacterium* sp. was found to be contaminating the *Chlorella vulgaris* culture with the likely species identified as *Microbacterium foliorum*, which is a common contaminant in most laboratory specimens (Behrendt et al., 2001), while *Pseudomonas tianjinensis* is cosmopolitan in distribution and can be found in soils, natural waters, as well as fertilised agricultural lands (Chen et al., 2018). Lastly, *Nannochloropsis limnetica* was contaminated by an *Agrobacterium* sp., with the best match being *Agrobacterium radiobacter* which is also commonly found in natural habitats; particularly around farmlands (Basavand et al., 2022). Depending on the strain, it could be pathogenic to plants. Here we provide a confident genus level identification of contaminants however we cannot be certain of the exact species identified through 16s sequencing alone. Further characterisation to be positive of the species identification would require deeper sequencing techniques such as using other gene markers or whole genome sequencing.

All identified microbes were gram-negative species, with *S. maltophilia, M. foliorum* and *S. chilensis* known to be multi-drug resistant.

## 4 Discussion

Photosensitisation has previously been demonstrated to be a good method for targeting metastasised carcinoma, as well as infections that are resistant to traditional antibiotics or chemical treatments. Cossu et al., 2016 demonstrated significant decrease in bacterial loads in wash waters with the use of the Rose Bengal dye. The mechanism associated to the observed antibacterial and antiviral effects upon exposure to light using Rose Bengal is attributed to the production of oxygen singlet radicals that result in non-specific cell-membrane damage. The phototrophic nature of the chosen microalgal species allow them to survive for extended periods in anoxic conditions so long as they have adequate access to light for photosynthesis, while the obligate aerobic bacteria were expected to be affected by the lack of oxygen. Based on the results from Table 1 and 2 however, it was evident that anoxic conditions; though detrimental to bacteria, also affected the microalgal population.

The use of Rose Bengal was thus investigated for its anti-bacterial effects. Based on the bacterial populations in oxic conditions noted in Tables 1 and 2, it was clear that as a standalone treatment, Rose Bengal was ineffective in removing bacterial contaminants within the chosen concentration range. However, in conjunction with anoxic conditions, it was able to synergistically remove bacterial contaminants without significantly affecting the microalgal population. The trend noted in Figure 1 indicates that ratio of bacteria to microalgae <1 was favourable as the bacterial population was lower than that of the microalgae. In all the species, anoxic conditions with Rose Bengal concentrations of 3 µM was found to produce ratios <1. Gram negative species have negatively charged extracellular membranes, making them particularly resistant to Rose Bengal (Waite × Yousef, 2009) which is anionic compared to Gram positive bacteria that are known to be susceptible to the compound even under dark conditions (Nakonechny et al., 2019).

On the contrary, none of the bacterial species noted in Table 3 have evolved to be facultative anaerobes and consequently do not have the ARC (Anoxic Redox Control) to cope with the change in oxic conditions. Dysfunctional ArcA mutants have been known to be susceptible to sodium dodecyl sulphate (SDS); a negatively charged detergent despite the membrane surface retaining its negative charge (Fu et al., 2015). Moreover, divalent ions such as Ca^2+^ and Mg^2+^ are known to promote Rose Bengal uptake in bacteria irrespective of Gram-staining status (George et al., 2009). This could explain the synergistic effect of Rose Bengal under anoxic conditions despite all the contaminant bacteria being Gram negative.

**Table 3:** Bacterial contaminants identified in the microalgal mother cultures using 16s sequencing.

| Culture | Genus identified | Top NCBI match | Percent identity |
| --- | --- | --- | --- |
| *Desmodesmus armatus* | *Stenotrophomonas* | *Stenotrophomonas maltophilia* | 99.67% |
|  | *Sphingopyxis* | *Sphingopyxis chilensis* | 97.88% |
| *Tetrademus obliquus* | *Stenotrophomonas* | *Stenotrophomonas maltophilia* | 98.17% |
|  | *Sphingopyxis* | *Sphingopyxis chilensis* | 98.69% |
| *Chlorella vulgaris* | *Microbacterium* | *Microbacterium foliorum* | 98.30% |
|  | *Pseudomonas* | *Pseudomonas tianjinensis* | 99.53% |
| *Nannochloropsis limnetica* | *Stenotrophomonas* | *Stenotrophomonas maltophilia* | 99.44% |
|  | *Agrobacterium* | *Agrobacterium radiobacter* | 99.50% |

Since no single method can unequivocally attest to axeny (Asatryan et al., 2022), the status was investigated by considering three pieces of evidence gathered from light microscopy, LB plate growth, and 16s PCR. Additional methods such as confocal microscopy (Sonowal et al., 2022), 16/18 s amplification (Vu et al., 2018), and electron microscopy (Asatryan et al., 2022; Vu et al., 2018) have been used by other researchers to attest for axeny.

### 4.1 Caveats and considerations

There are significant considerations when applying anoxic Rose Bengal to achieve axenic cultures; many aspects that could not be addressed within the experimental setup described herein. Firstly, all the contaminating microbes identified (Table 3) were obligate aerobes. Thus, the anoxic conditions were able to significantly affect their populations within the mother culture. Moreover, the cellular uptake required for the synergistic antimicrobial action of Rose Bengal despite the bacteria being gram-positive, was made possible only by the uptake enhanced by the Ca^2+^ and Mg^2+^ ions that are generally available in higher concentrations in BBM compared to natural habitats to promote microalgal growth. The effectiveness of the described treatment is unclear if facultatively anaerobic microbes were to contaminate cultures under low Ca^2+^ and Mg^2+^ concentrations.

Secondly, only bacterial contamination was addressed in this work as fungal, amoebic, or other contaminating lifeforms were not encountered within the mother cultures and were consequently not considered as part of this investigation. The effectiveness of the anoxic Rose Bengal in the proposed 3 µM concentration remains unclear under such contaminating conditions.

Lastly, this method was designed to only serve in unialgal cultures; i.e., only one microalgal species in a given mother culture. Although different microalgae showed different degrees of viability to the anoxic Rose Bengal treatment (Table 2), and this may be leveraged to obtain pure cultures of the more resistant microalgae, physical methods such as micropicking, microfluidics or flow-cytometry may be a more suitable intermediate step to separate individual microalgal species.

## 5 Conclusion

Decontamination appears to be largely a function of anoxy that is synergistically enhanced by Rose Bengal at concentrations ≥3 µM. Most cultures were contaminated by Gram negative bacteria, a couple of which were multi-drug resistant and consequently, antibiotic treatment was ineffective in decontaminating the cultures.

## Acknowledgements

A.I is a grateful recipient of the Irish Research Council (IRC) Postdoctoral Fellowship (GOIPD/2023/1240), and UCD College of Engineering × Architecture (CEA) seed fund. M.M is thankful to the Erasmus mobility award that facilitated his travel to UCD.

